# Propylene glycol-based antifreeze as an effective preservative for DNA metabarcoding of benthic arthropods

**DOI:** 10.1101/2020.02.28.970475

**Authors:** Chloe V. Robinson, Teresita M. Porter, Michael T.G. Wright, Mehrdad Hajibabaei

## Abstract

Maintaining the integrity of DNA in bulk environmental samples from source to laboratory is crucial for capturing the true range of taxa present within an ecosystem. Preservation consideration of DNA is particularly important if samples are being collected in remote areas and by non-specialist ‘citizen scientists’ in nationwide programs. Traditionally, absolute ethanol is used as the preferred preservative for environmental samples collected for downstream DNA metabarcoding analyses. However, transport, shipping and DNA extraction of samples preserved in ethanol is a lengthy procedure due to safety restrictions and the requirement of full ethanol evaporation prior to extraction. We examined the efficacy of an easily accessible, non-toxic propylene glycol-based antifreeze as an alternative to absolute ethanol for preserving macroinvertebrate DNA from bulk-benthos DNA samples. We tested the differences in both cytochrome oxidase I (COI) exact sequence variants (ESVs) and COI taxonomic orders detected in both ethanol and antifreeze samples using two processing methods (no evaporation of preservative versus full evaporation). In addition, we assessed the detection of families and genera within the Arthropoda phylum for preservative type, site and processing method. Our results suggest that antifreeze is a suitable alternative to ethanol, a greater global ESV richness reported for antifreeze samples. Additionally, a higher proportion of arthropod reads in ESVs were detected in antifreeze (average 69%) compared with ethanol (average 53%). Finally, antifreeze samples produced similar results for the different processing methods, whereas ethanol samples failed to produce similar results without prior evaporation. Although ethanol is currently widely used for DNA preservation, our results demonstrate that by using antifreeze, it is possible to achieve similar taxonomic coverage and community assemblages of bulk-benthos DNA samples for macroinvertebrates, with the added simplicity and shorter laboratory processing time achieved using an easily available, unregulated preservative.

## Introduction

Increasingly, DNA-based techniques are being used to detect and monitor biodiversity across a variety of ecosystems (Baird & Hajibabaei, 2012; Taberlet, Coissac, Hajibabaei, & Rieseberg, 2012; Taberlet, Bonin, Zinger, & Coissac, 2018; Thomsen & Willerslev, 2015; M Hajibabaei, Baird Donald J., Fahner Nicole A., Beiko Robert, & Golding G. Brian, 2016). A prominent approach, DNA metabarcoding, involves sequencing taxonomic marker genes such as DNA barcodes from bulk environmental samples such as benthos (Taberlet et al., 2012; Goldberg, Strickler, & Pilliod, 2015; Pedersen, Mikkel Winther et al., 2015; Thomsen & Willerslev, 2015; Minamoto, Naka, Moji, & Maruyama, 2016). Effective preservatives are required to prevent DNA degradation of environmental samples, and depending on the type of sample and target taxa, optimal DNA preservation methods will vary (Sales, Wangensteen, Carvalho, & Mariani, n.d.; Barnes & Turner, 2016; Koziol et al., 2019). Maintaining the viability of DNA from source to laboratory is particularly important for research involving an element of community-based monitoring (i.e. citizen science) in the sampling process (Steininger, Storer, Hulcr, & Lucky, 2015). Often, the biggest concern with citizen science biomonitoring initiatives is the quality and consistency of data collected (Kelling et al., 2015; Kosmala, Wiggins, Swanson, & Simmons, 2016), especially with the additional concern of DNA contamination (Biggs et al., 2015; Dickie et al., 2018). Cross-contamination of DNA can occur between samples and sites, which can produce false positives and is highly problematic when attempting to infer community patterns through metabarcoding-based biodiversity assessments (Dickie et al., 2018; Taberlet et al., 2018; Zinger et al., 2019).

Environmental DNA metabarcoding enables rapid assessment of ecosystem health, primarily through assessing biodiversity patterns of bioindicator species such as freshwater benthic macroinvertebrates (Mehrdad Hajibabaei, Spall, Shokralla, & van Konynenburg, 2012; M Hajibabaei et al., 2016). The simple, quick and easily repeatable sampling protocol associated with freshwater sampling provides a key opportunity to integrate community-based efforts with sample collection (Biggs et al., 2015; Buxton et al., 2018; Robinson et al., 2018). Similar to sampling protocol simplicity, the DNA preservative of choice for community-based biomonitoring initiatives needs to be effective, easy to handle and uncomplicated to transport and/or store. Highly concentrated ethanol (>99%) is a known effective preservative for DNA from a range of taxa and environment types (Mehrdad Hajibabaei et al., 2012; Steininger et al., 2015) and is widely utilised to preserve eDNA and bulk tissue samples (Sales et al., n.d.; K. E. Williams, Huyvaert, & Piaggio, 2016; Minamoto et al., 2016; Koziol et al., 2019). Ethanol however is classified as a ‘dangerous good’, meaning complicated and costly arrangements for storage and transit of environmental samples preserved in ethanol, particularly regarding air transportation (S. T. Williams, 2007; Steininger et al., 2015). This is problematic when attempting to collect environmental samples from remote locations (i.e. air access only), which in turn directly limits the geographical extent of DNA-based biomonitoring (McGee, Robinson, & Hajibabaei, 2019). In addition, samples preserved in ethanol require a lengthy evaporation period prior to DNA extraction, due to the PCR inhibitory nature of ethanol in concentrations above 1% (Demeke & Jenkins, 2010; Schrader, Schielke, Ellerbroek, & Johne, 2012). This substantially increases the laboratory processing time of ethanol-preserved environmental samples (Demeke & Jenkins, 2010; Schrader et al., 2012). Another issue with relying on ethanol as a preservative, is the current by-laws regarding prohibition of alcohol in First Nation communities across countries including Canada (Brady, 2000; Campbell, 2016). These issues overall can result in limits to the freshwater locations which can be assessed for biodiversity using DNA metabarcoding, which ultimately results in large data deficiencies (Pereira & David Cooper, 2006).

Across Canada, a large proportion of watersheds are currently classified as ‘Data Deficient’ (WWF-Canada, 2017). Out of 167 sub-watersheds, currently 110 lack the data necessary to determine baseline health status (WWF-Canada, 2017). In terms of benthic invertebrates, 112 sub-watersheds are data deficient, and only 35 sub-watersheds have good to very good benthic invertebrate health (remaining 20 fall below this threshold; (WWF-Canada, 2017). In order to combat this data deficiency, the STREAM (Sequencing The Rivers for Environmental Assessment and Monitoring) project was established with the main aim of combining community-based monitoring with DNA metabarcoding to assess the benthic macroinvertebrate communities in watersheds (particularly with data deficient status) across Canada.

The collaborative approach of STREAM combines academic (University of Guelph) and non-academic partners (World Wildlife Fund Canada (WWF-Canada), Living Lakes Canada (LLC) and Environmental and Climate Change Canada (ECCC)), to facilitate citizen training and genomic analyses to create a baseline of watershed health. For this project, a number of First Nation groups are to participate in sample collection and some target watersheds and sub-watersheds exist in remote locations; to achieve the desired sample spatial coverage and community involvement, an effective, safe alternative to ethanol (and other conventional methods such as freezing) needs to be determined.

Previously, a study compared the DNA preserving ability of easily accessible glycol-based antifreeze with ethanol for short-term preservation of invertebrate DNA and determined that antifreeze was equally as effective at preserving DNA, even after seven days under direct outdoor exposure (Steininger et al., 2015). Similarly, other studies have investigated the relative suitability of a range of other preservatives on environmental samples, including sodium acetate-ethanol solution (Ladell, Walleser, McCalla, Erickson, & Amberg, 2019), Longmire’s solution (K. E. Williams et al., 2016) and cationic surfactant (Yamanaka et al., 2017), however these have focused on the preservation of water eDNA as opposed to DNA derived from benthos.

The aim of this study therefore was to compare the efficiency of commonly used molecular grade ethanol (>99%) with commercially available propylene glycol-based antifreeze (Absolute Zëro RV Waterline Antifreeze; Recochem, Montreal QC), to investigate the taxonomic coverage of benthic macroinvertebrate communities, in particular arthropods, detected through next-generation sequencing (NGS) of benthic samples, using two laboratory processing methods (no evaporation of preservative prior to DNA extraction and full evaporation).

## Materials and Methods

### Sample design and study sites

In order to accurately assess any differences in arthropod detection using 99% EtOH or antifreeze, firstly, single (‘mock’) benthic aquatic invertebrate community (hereafter designated as ‘benthos’) samples were taken from riffle locations at three sites within tributaries of the Grand River (Waterloo, ON) (Table S1). These samples were homogenised prior to preservative addition, to reduce the influence of natural variation of arthropods masking the effect of the treatment (preservative type used). In addition, paired benthos samples were collected from three additional sites and preserved prior to homogenisation.

All samples were collected using a standard Canadian Aquatic Biomonitoring Network (CABIN) kick net with a 400μm mesh net and attached to a pole and net frame. Effort was standardized to a 10m transect collected over three minutes per sample (Environment and Climate Change Canada, 2012). After collecting samples, nets were drained, the benthic material was removed from the net and placed in a sterile white 1L polyethylene sample jar filled no more than half full. The net was rinsed and inspected to remove any remaining invertebrates. To avoid DNA carryover between sites, sterile nets were used to collect samples at each site and field crew wore clean nitrile gloves to collect and handle samples in the field and laboratory. Samples (n = 9), were placed on ice in a cooler for transport to the laboratory the same day.

Once at the laboratory, 99% EtOH was added to one replicate from the three sites with paired samples and Absolute Zëro RV Waterline Antifreeze was added to the remaining replicate, at a ratio of 1:3 preservative to sample.

### Sample homogenisation and DNA extraction

For the single ‘mock’ samples, a crude homogenate was produced by adding 300 mL molecular grade water and blending the component of each sample using a standard blender that had been previously decontaminated and sterilized using ELIMINase™ followed by a rinse with deionized water and UV treatment for 30 min. After blending, the homogenate was split into two sample jars and EtOH was added to one sample jar and antifreeze to the remaining jar. These samples remained on the laboratory bench at room temperature for 3 days. For the paired samples, a crude homogenate was produced following the steps above and samples of each homogenate from paired and single samples was then transferred to individual 50 mL Falcon tubes and centrifuged at 2400 rpm for 2 minutes to pellet the tissue. The supernatant was removed, and at this point, two methods of sample processing were tested. For each antifreeze sample and for three ethanol samples, one subsample of homogenate was extracted without evaporating excess preservative from the pellet (Method 1) and one subsample was extracted after excess preservative was fully evaporated from the pellet (Method 2). Only three ethanol samples were selected as protocols have already established the inhibiting effect of ethanol on DNA extraction, therefore these samples were controls. Subsamples which had preservative evaporated were dried at 70°C, until the preservative was fully evaporated (approximately 4 - 8hr). Using a sterile spatula, ~300 mg dry weight of homogenate was subsampled into PowerBead tubes containing garnet. The remaining dry mass was stored in the Falcon tubes at −20°C as a voucher.

DNA was extracted using a DNeasy PowerSoil kit (Qiagen, CA) following manufacturer’s instructions. DNA was eluted in a final volume of 50uL of buffer C6. Purity and concentration of DNA for each site was checked using a NanoDrop spectrophotometer and recorded. Samples were kept at −20°C for further PCR and sequencing. Each batch of DNA extractions included a negative control where no tissue was included. 50mL of each of the source ethanol and antifreeze were filtered and extracted using Qiagen’s DNeasy PowerWater kit according manufacturer’s protocol to verify the purity of the source preservatives.

### Library preparation and high-throughput sequencing

Three fragments within the standard COI DNA barcode region were amplified with the following primer sets (B/ArR5 [~310 bp] called BR5, LCO1490/230_R [~230 bp] called F230R and mICOIintF/jgHCO2198 [~313 bp] called ml-jg (Mehrdad Hajibabaei et al., 2012; Geller, Meyer, Parker, & Hawk, 2013; Leray et al., 2013; Gibson et al., 2014)) using a two-step PCR amplification regime. The first PCR used COI specific primers and the second PCR involved Illumina-tailed primers. The PCR reactions were assembled in 25 μl volumes. Each reaction contained 2 μl DNA template, 17.5 μl molecular biology grade water, 2.5 μl 10× reaction buffer (200 mM Tris-HCl, 500 mM KCl, pH 8.4), 1 μl MgCl2 (50 mM), 0.5 μl dNTPs mix (10 mM), 0.5 μl forward primer (10 mM), 0.5 μl reverse primer (10 mM), and 0.5 μl Invitrogen’s Platinum Taq polymerase (5 U). The PCR conditions were initiated with heated lid at 95°C for 5 min, followed by a total of 35 cycles of 94°C for 40 s, 46°C for 1 min, and 72°C for 30 s, and a final extension at 72°C for 5 min, and hold at 4°C. The amplicons from the first PCR were used as templates in the second PCR with the same amplification condition used in the first PCR with the exception of using Illumina-tailed primers in a 35-cycle amplification regime. All PCRs were done using Eppendorf Mastercycler ep gradient S thermalcyclers. Negative control reactions (no DNA template) for antifreeze, ethanol, DNA extraction kit and for the PCRs were included in all experiments.

PCR products were visualized on a 1.5% agarose gel to check the amplification success. Amplicons were purified using MinElute PCR Purification kit (Qiagen) eluting with 15uL molecular grade water. Purified 2^nd^ PCR amplicon samples were quantified using a QuantIT PicoGreen dsDNA assay kit and normalized to the same concentration based on those values. Samples were then dual indexed using Illumina’s Nextera XT indexes (FC-131-1002) and pooled into a single tube. The pooled library was purified using AMpure magnetic beads, quantified using a QuantIT PicoGreen dsDNA assay, and fragment length was determined using a Bioanalyzer. The purified library was diluted based on the concentration and average fragment length and sequenced on an Illumina MiSeq using a V3 MiSeq sequencing kit (300 × 2; AAS-102-2003), with a PhiX spike-in of 10%.

### Bioinformatic processing

Raw Illumina paired-end reads were processed using the SCVUC v4 pipeline available from https://github.com/Hajibabaei-Lab/SCVUC_COI_metabarcode_pipeline. Briefly, raw reads were paired with SeqPrep v1.3.2 ensuring a minimum Phred score of 20 and minimum overlap of at least 25 bp (St. John, J, 2016). Primers were trimmed with CUTADAPT v2.6 ensuring a minimum trimmed fragment length of at least 150 bp, a minimum Phred score of 20 at the ends, and allowing a maximum of 3 N’s (Martin, 2011). All primer-trimmed reads were concatenated for a global ESV analysis. Reads were dereplicated with VSEARCH v2.14.1 using the ‘derep_fulllength’ command and the ‘sizein’ and ‘sizeout’ options (Rognes, Flouri, Nichols, Quince, & Mahé, 2016). Denoising was performed using the unoise3 algorithm implementation in VSEARCH with the ‘cluster_unoise’ command (Edgar, 2016). This method removes sequences with potential errors, PhiX carry-over from Illumina sequencing, and rare reads. Here we defined rare reads to be exact sequence variants (ESVs) containing only 1 or 2 reads (singletons and doubletons; (Callahan, McMurdie, & Holmes, 2017)). Putative chimeric reads were removed in a separate step using the uchime3 algorithm implementation in VSEARCH using the ‘uchime3_denovo’ command. Putative pseudogenes were identified and removed by translating denoised ESVs into every possible reading frame (plus strand) using ORFfinder v0.4.3 (NCBI Resource Coordinators, 2018) keeping the longest open reading frame (ORF), calculating cutoffs to identify ORFs with outlier lengths (shorter than the 25^th^ percentile - 1.5 * interquartile length or longer than the 75^th^ percentile + 1.5 * interquartile length). We used the mitochondrial invertebrate genetic code for translation, so all subsequent analyses were limited to arthropoda. An ESV x sample table was created with VSEARCH using the ‘usearch_global’ command, mapping reads to ESVs with 100% identity. ESVs were taxonomically assigned using the COI Classifier v4 that uses the RDP classifier naive Bayesian classifier v2.12 (Wang, Garrity, Tiedje, & Cole, 2007).

### Statistical analyses

Diversity analyses were conducted in Rstudio v 1.1.456 using R v 3.5.1 (RStudio Team, 2016) with the vegan package (Dixon, 2003; RStudio Team, 2016). To assess sequencing depth for both preservative types, we plotted rarefaction curves using a vegan ‘rarecurve’ function. Unless otherwise stated, read depth per sample was normalized to the 15^th^ percentile library size using the ‘rrarefy’ function in vegan. We assessed the recovery of ESVs from ethanol samples compared with antifreeze samples and assessed the proportion of all ESVs that could be taxonomically assigned with high confidence. Taxonomic assignments were deemed to have high confidence if they had the following bootstrap support cutoffs: species >= 0.70 (95% correct), genus >= 0.30 (99% correct), family >= 0.20 (99% correct) as is recommended for 200 bp fragments using the COI Classifier (Porter & Hajibabaei, 2018). Assignments to more inclusive ranks, ex. order, do not require a bootstrap support cutoff to ensure that 99% of assignments are correct.

We calculated ESV richness across different groups of the data to compare differences across sites and preservation methods (ethanol or antifreeze samples). To check for significant differences, we first checked for normality using visual methods (ggdensity and ggqqplot functions in R) and the Shapiro-Wilk test for normality (Shapiro & Wilk, 1965).

For assessing overall community structure of arthropods (from post-evaporation data only) detected from different preservatives, we used non-metric multi-dimensional scaling analysis (NMDS) on Sorensen dissimilarities (binary Bray-Curtis) using the vegan ‘metaMDS’ function. A Shephard’s curve and goodness of fit calculations were calculated using the vegan ‘stressplot’ and ‘goodness’ functions and to assess the significance of groupings, we used the vegan ‘vegdist’ function to create a Sorensen dissimilarity matrix, the ‘betadisper’ function to check for heterogeneous distribution of dissimilarities, and the ‘adonis’ function to do a permutational analysis of variance (PERMANOVA) to check for any significant interactions between groups (preservative, site, processing method). To maintain a balanced design during statistical testing, we limited comparisons to post-evaporation samples.

To investigate arthropod ESVs and orders shared between preservative types, sites and processing methods, we used the VennDiagram function ‘draw.venn’ in R. We also plotted the number of arthropod ESVs detected from ethanol or antifreeze preserved samples, limited to post-evaporation samples for a balanced comparison. We calculated the Jaccard index to look at the overall similarity in arthropod community structure between ethanol and antifreeze samples. We also visualized the frequency of ESVs detected from arthropod families using a heatmap generated using geom_tile (ggplot) in R, limited to post-evaporation samples for a balanced comparison.

## Results

Antifreeze samples produced clear bands in the gel for both processing methods for both PCR reactions (Figure S2). Ethanol samples generated bands similar to antifreeze samples for all sites. A total of 11,185,532 x2 Illumina paired-end reads were sequenced (Table S2). After bioinformatic processing, we retained a 19,298 ESVs (9,908,826 reads). After taxonomic assignment, a total of 5,198 arthropod ESVs (2,418,452 reads) were retained for data analysis (Table S3).

Out of all ESVs, 34.8% were assigned to Arthropoda, accounting for 60.5% of reads in all ESVs (Figure S3). Rarefaction curves that reach a plateau show that our sequencing depth was sufficient to capture the ESV diversity in our PCRs (Figure S4). In terms of classification, most arthropod genera and families and a majority of arthropod species were confidently identified (Figure S5).

Negative controls (antifreeze, ethanol, extraction kit and PCR), produced a small number of sequences from a total of 13 orders (Table S4), however the low number of ESVs returned (between 3 and 80) would not have influenced the sequencing results observed in the field samples dataset (Hornung, Zwittink, & Kuijper, 2019).

### Taxonomic coverage

For the single ‘mock’ samples, alpha diversity calculated from mean overall ESV richness, displayed very similar values for antifreeze (229) and ethanol (222) samples (after normalization). For paired samples, alpha diversity was again similar between antifreeze (251) and ethanol (245) samples. In terms of the two different processing methods for the paired samples, method 1 (no evaporation prior to extraction) resulted in marginally higher ESV richness for antifreeze (273 method 1 and 271 method 2) and as expected, ethanol samples produced greater ESV richness post-evaporation (196 method 1 and 220 method 2).

### Arthropod Diversity

To address whether antifreeze could be effective at preserving arthropod taxa, we investigated results obtained for the Arthropoda phylum. Arthropod ESV richness was relatively similar between ethanol and antifreeze samples post-evaporation (method 2), with replicates clustering close together for a majority of sites (Figure 1). PERMANOVA on post-evaporation samples (for a balanced design), shows that analyzing samples from paired samples or a single split sample (experiment) explains 19% of the variation in Bray Curtis dissimilarities (p-value = 0.001); storing samples in ethanol or antifreeze (treatment) explains 4% of the variation (p-value = 0.001); and that the interaction between experiment and treatment explains 4% of the variation (p-value = 0.001). We were not able to compare samples extracted before and after evaporation because we did not have a balanced design (not enough ethanol pre-evaporation ethanol, see Figure 1).

**Figure 1.**
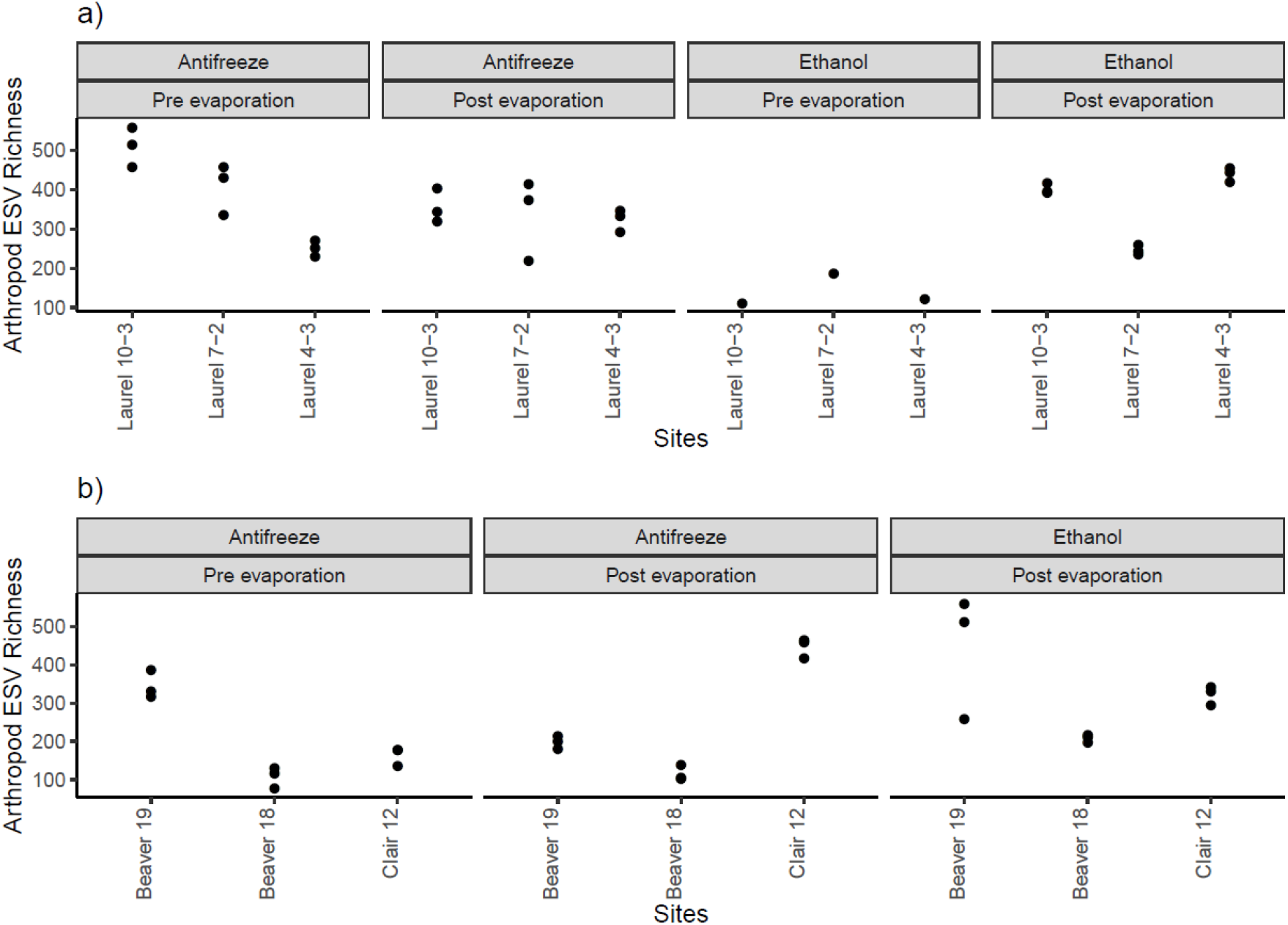
Arthropod ESV richness varies across pre- and post-evaporation antifreeze and ethanol stored samples. Results from two experiments based on a) paired sampling (two benthic samples collected nearby, each stored in antifreeze or ethanol) and b) a single sample split in two (then each half sampling stored in antifreeze or ethanol). Pre- and post-evaporation refers to benthic samples removed from storage solution (antifreeze or ethanol) and extracted before and after allowing liquid to evaporate before proceeding to DNA extraction. Based on rarefied data.

The Jaccard index for both the mock and paired sample datasets (analyzed independently) between preservative types (post-evaporation for balanced design) is 0.94, indicating samples are 94% similar. Within each preservative tested, method 1 and 2 resulted in the same arthropod ESV richness for antifreeze samples (231), however ethanol samples produced a greater arthropod ESV richness for method 2 (180 pre- and 193-post evaporation).

At the arthropod order level, there was a total of 22 orders shared between the two preservative types for single samples and 26 shared in the paired samples. Within the single samples, two orders were unique to ethanol (Mantodea and Siphonaptera) and four orders unique to antifreeze (Entomobryomorpha, Harpacticoida, Ixodida, Psocoptera), whereas the paired samples indicated four orders unique to ethanol and five unique to antifreeze (Figure 3). For site, 14 (single samples) and 13 (paired samples) arthropod orders were shared between all sites, with the most unique arthropod orders found at Clair 12 (4) and Laurel 10 (3). When looking at processing methods for antifreeze, 23 (single samples) and 27 (paired samples) arthropod orders were shared (Figure 3).

**Figure 2.**
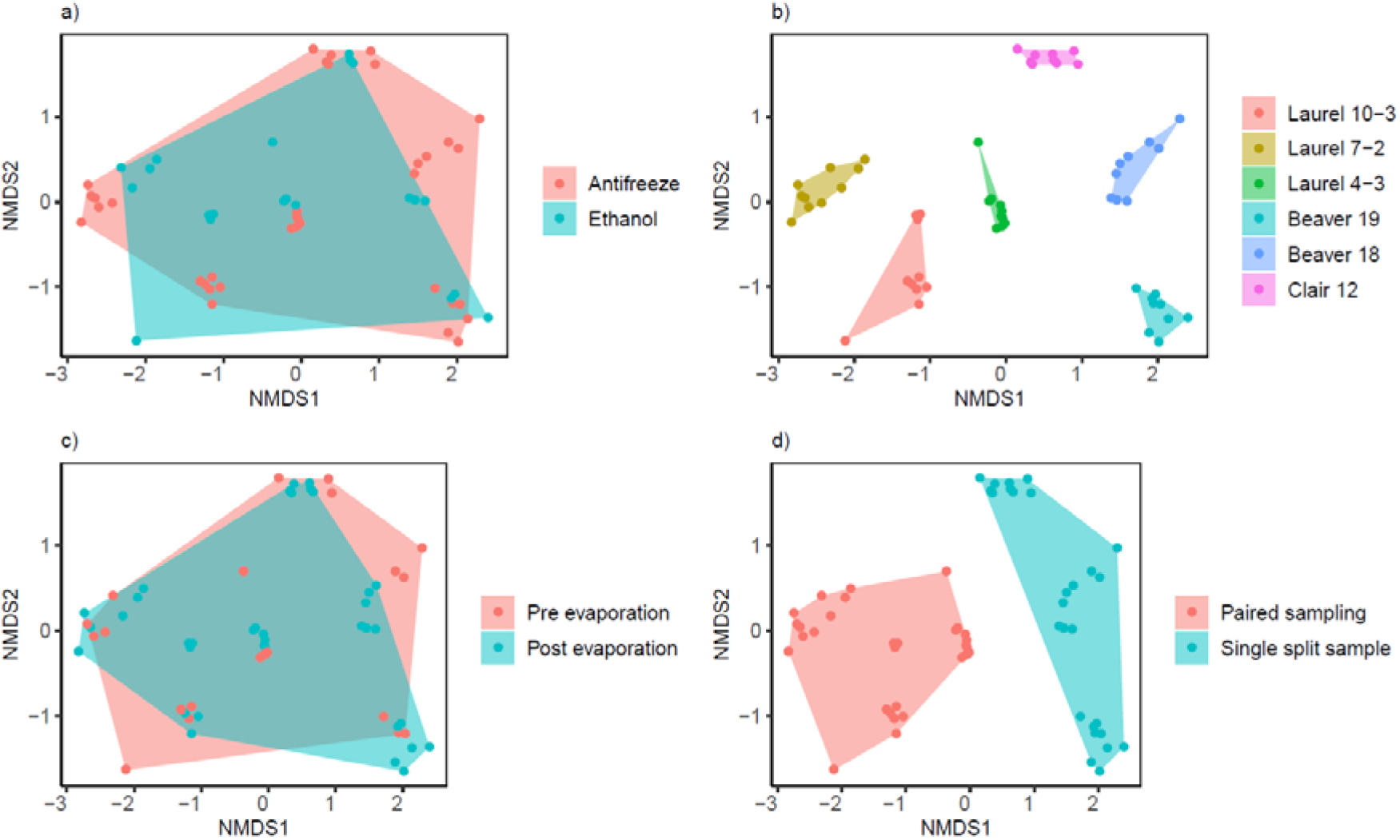
Arthropod samples cluster mainly by site and sampling method. Non-metric multi-dimensional scaling plots show a) overlap between the binary Bray Curtis (Sorensen) dissimilarities across samples stored in antifreeze and ethanol, b) clustering by sampling site, c) overlap between samples extracted before and after evaporation, and d) clustering based on field sampling method (stress = 0.124, R2 = 0.93). Based on rarefied data.

**Figure 3.**
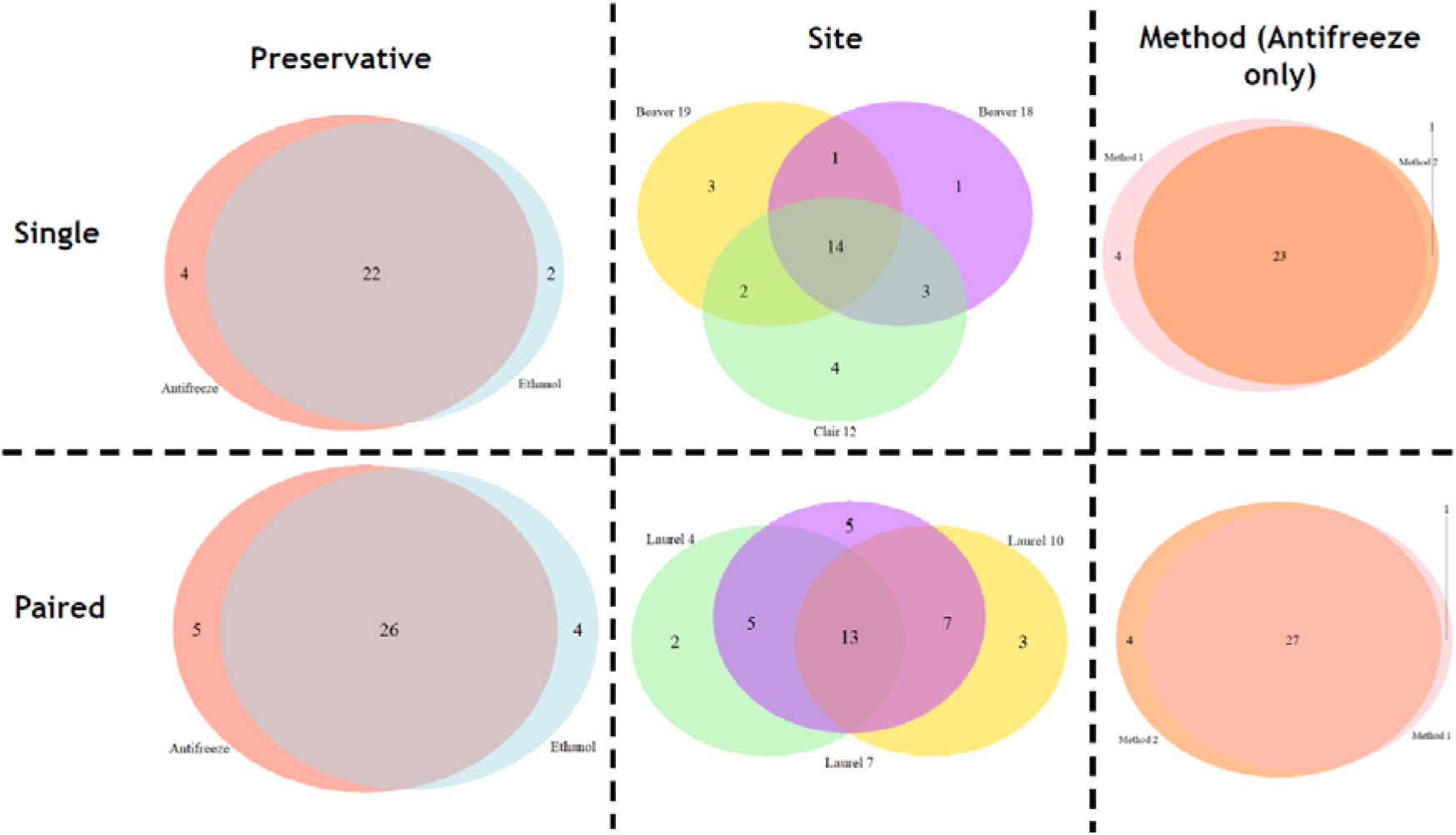
High degree of taxonomic overlap between treatment groups. Venn diagrams of unique and overlapping arthropod orders for preservative types, sites and evaporation method. Based on rarefied post-evaporation data for preservative and site.

Through investigating families from arthropod orders, a similar diversity of families was detected from antifreeze samples (Figure 4). Some of the confidently identified arthropod genera represented by more than 2(Log10) sequence variants, identified from ethanol and antifreeze, included: Caenis (Ephemeroptera), Polypedilum (Chironomidae), Tipula (Chironomidae) and Stenelmis (Coleoptera; Figure 4). In terms of the bioindicator orders and families for both the single and paired samples, the heatmap shows that antifreeze and ethanol samples showed similar results for the detected Ephemeroptera, Trichoptera and Plecoptera families across replicates at each site (Figure 5). Subtle differences in the detection between preservative types was evident in the single samples; in some cases, antifreeze samples produced a greater number of reads for families detected (e.g. Uenoidae; Figure 5b) and similarly in other cases, ethanol samples produced a greater number of reads for particular families detected (e.g. Leptophlebiidae; Figure 5b). A large majority of the families detected were present in both ethanol and antifreeze samples to the same read number extent across all sites.

**Figure 4.**
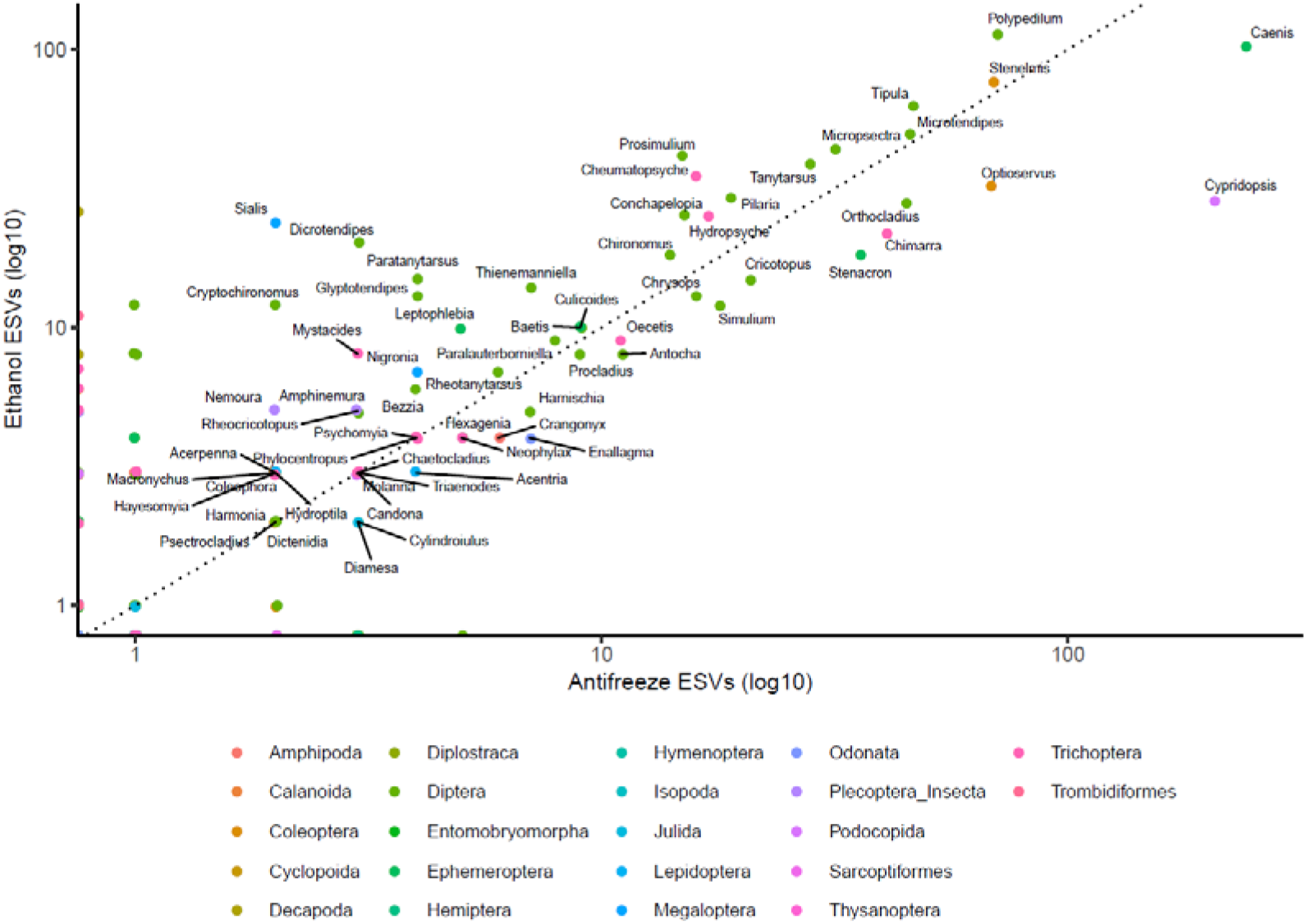
The number of arthropod ESVs detected from genera detected from samples preserved in antifreeze or ethanol are similar. Each point represents a genus identified with high confidence. The points are colour coded for the 22 arthropod orders detected in this study. A 1:1 correspondence line (dotted) is also shown. A log10 scale is shown on each axis to improve the spread of points with small values. Based on rarefied data. Only post-evaporation samples are shown for a balanced comparison.

**Figure 5.**
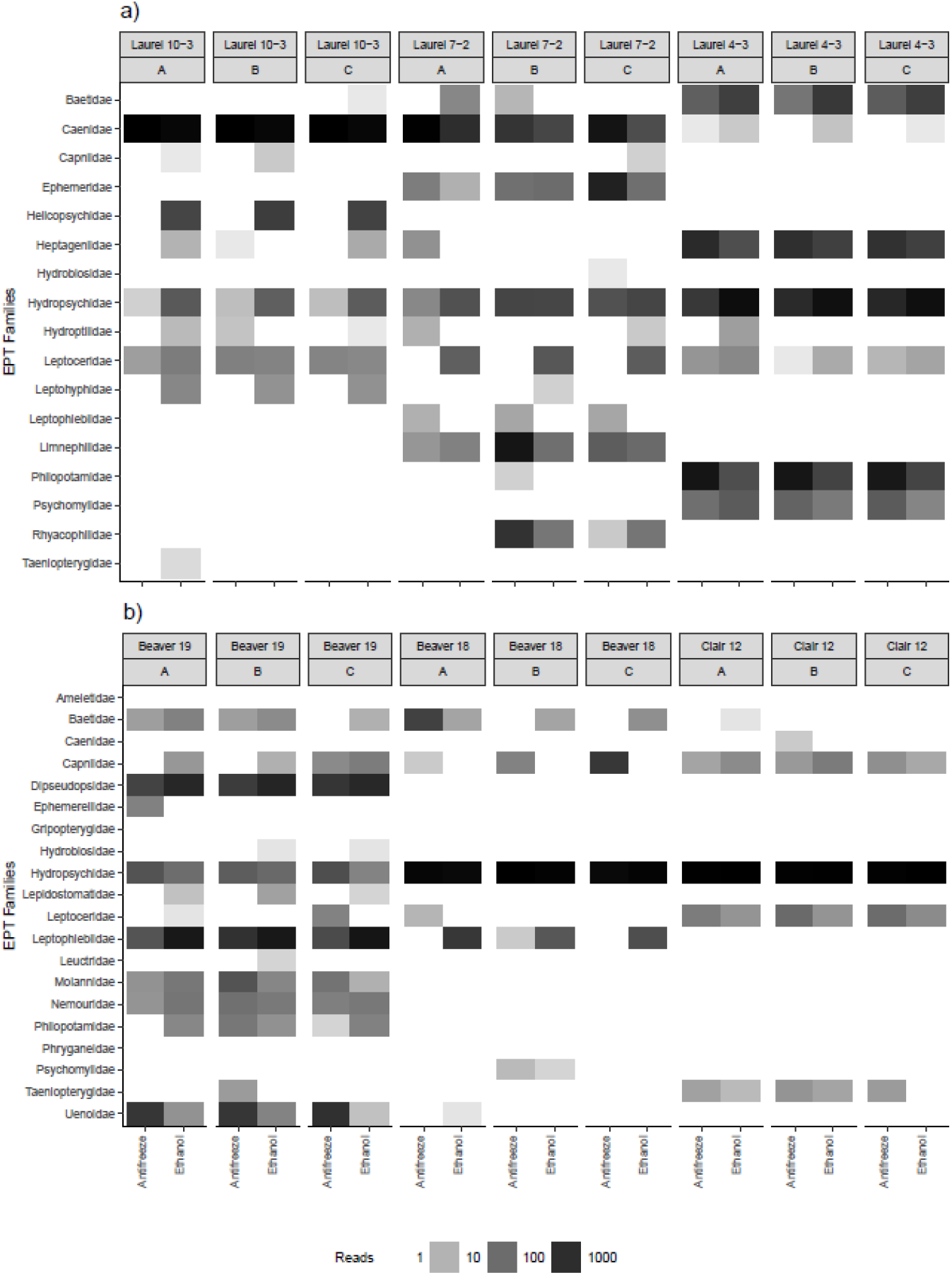
Replicates detect similar bioindicator families across experiments and treatments. Only ESVs taxonomically assigned to families with high confidence (bootstrap support >= 0.20) are included. Part a) shows a paired sampling design, and b) a single sample split into two parts for each treatment. For each site, three replicates (A, B, C) are shown for each treatment (storage in antifreeze or ethanol). The heatmap shows the detection of Ephemeroptera, Trichoptera, or Plecoptera (EPT) families across samples. Based on normalized data. Only post-evaporation samples are shown.

## Discussion

For monitoring of biodiversity, it is vital that DNA preservatives used in metabarcoding studies are effective at preventing DNA degradation, to enable the capture of taxa present within an environmental sample (Rees, Maddison, Middleditch, Patmore, & Gough, 2014; Strickler, Fremier, & Goldberg, 2015; Barnes & Turner, 2016). Commonly, 95-100% ethanol is the standard preservative used for maintaining the integrity of both tissue and environmental samples containing DNA, including freshwater, seawater and bulk-benthos (Baird & Hajibabaei, 2012; Mehrdad Hajibabaei et al., 2012; Rees et al., 2014). However, this ‘one size fits all’ approach is unlikely to be feasible for optimal DNA preservation across all environment types, due to existing variation in taxa and abiotic factors (Barnes & Turner, 2016). Typically, bulk-samples contain less degraded DNA than eDNA (i.e. free DNA existing in a water medium), and therefore often a wider coverage of biodiversity is detected using this sample type (Taberlet et al., 2012; Turner, Uy, & Everhart, 2015; Mehrdad Hajibabaei et al., 2019). Additionally, biomonitoring research should strive to sample taxa from a wide geographic range, including remote locations, to prevent a location bias and therefore potential under-/over-representation of taxa (McGee et al., 2019). Metabarcoding bulk-benthos single ‘mock’ and paired samples provided an opportunity to test the effectiveness of a proposed alternative preservative to ethanol for preserving macroinvertebrate DNA. Our study has shown that arthropod ESV richness is comparable between conventional absolute ethanol and commercial antifreeze (Absolute Zëro RV Waterline Antifreeze) and that arthropod community assemblages are highly similar for both types of preservative.

To determine any true effect of preservative type used on detection of macroinvertebrate DNA, the approach of using single ‘mock’ samples was important, as natural biological variation can be high even within the same site, as demonstrated in Figure 1 and 5. We expected to observe higher levels of dissimilarity between ethanol and antifreeze samples in the paired samples compared to the single samples. However, overall ESV richness remained similar and for arthropods, similarity of community assemblage remained at 94% between the two preservative types for paired samples. Preservatives used to maintain the integrity of DNA for biomonitoring are required to be consistent, in terms of preserving various taxa across different sites, to enable an accurate assessment of existing community assemblages. We have demonstrated in this study that antifreeze has the capacity to preserve macroinvertebrates, in particularly benthic arthropods, consistently at the same capacity as conventional ethanol.

Unlike ethanol, it is apparent that antifreeze does not appear to inhibit the DNA extraction or PCR in the processing of samples. In terms of laboratory processing, optimizing DNA extraction protocols to generate results faster whilst still maintaining high quality and reliable results is key for reducing consumable and labor costs for biomonitoring research (Bonada, Prat, Resh, & Statzner, 2006; Baird & Hajibabaei, 2012; Vasselon, Domaizon, Rimet, Kahlert, & Bouchez, 2017). For biomonitoring studies, the shift from morphological taxonomic identification to DNA metabarcoding has already significantly cut costs of downstream analyses (Baird & Hajibabaei, 2012; Muha, Rodríguez-Rey, Rolla, & Tricarico, 2017). However, DNA metabarcoding research can be limited by PCR inhibition, and often time-consuming steps are taken in order to fully remove the ethanol preservative and purify DNA prior to sequencing (Demeke & Jenkins, 2010; Schrader et al., 2012). Using propylene-glycol based antifreeze as a preservative for environmental samples, allows for the lengthy preservative evaporation to be bypassed and instead DNA pellets formed from centrifuged samples can be extracted directly and instantly. In terms of the STREAM project, the application of antifreeze as the preservative of choice will greatly facilitate the timely turnaround of results.

For many freshwater biomonitoring programmes, the main aim is to investigate and compare macroinvertebrate community assemblages across sites of differing geographic and quality status, to build a picture of freshwater ecosystem health. Results from this study have shown that despite the use of two different preservatives using two processing methods, the main source of variation in the data was derived from the sample design (single or paired). Despite the treatment (preservative used) significantly explaining 4% of the variation, there was an overlap of samples by treatment and samples clustered more by sampling site. It is common that different microhabitats are located within rivers and that small variations in environmental conditions such as pH, dissolved oxygen and organic compounds coupled with anthropogenic use can alter macroinvertebrate assemblages, particularly of bioindicator species, on a local scale (Mehrdad Hajibabaei et al., 2012; Emilson et al., 2017; Pawlowski et al., 2018; Yang, Jeppe, Pettigrove, & Zhang, 2018). Moreover, most arthropod orders were shared across ethanol and antifreeze samples, and we have shown that the traditional bioindicator orders being shared among the preservative types, demonstrating the similar performance of antifreeze to traditionally used ethanol.

Overall, antifreeze performed similarly to ethanol as a preservative of macroinvertebrate DNA. Comparable recovery of arthropod ESVs and matched detection of key bioindicator orders and families in antifreeze samples highlights the suitability of propylene glycol-based antifreeze as an alternative to conventional absolute ethanol. Use of inexpensive, easily accessible antifreeze, such as the brand used in this study, will greatly benefit DNA-based citizen science initiatives, in terms of feasibility for sampling remote sites, sample preservation, laboratory processing time, and simplifying the sample shipping process. Additional studies into the long-term storage of DNA in antifreeze and applicability of antifreeze for additional environmental sample types, such as water and soil, would be beneficial to further investigate the potential of propylene glycol-based antifreeze as a widely used DNA preservative.

**Table 1.**
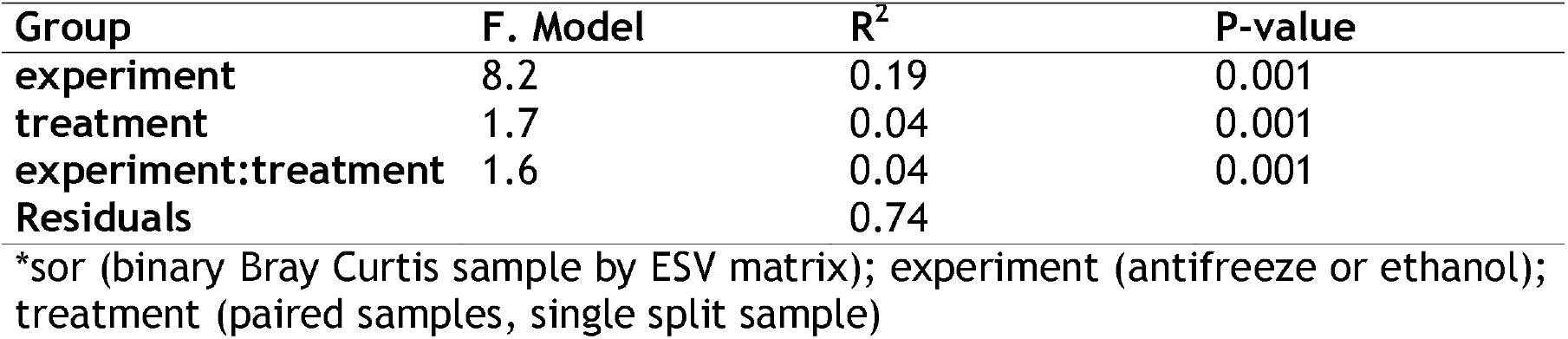
PERMANOVA formula sor ~ experiment * treatment (strata = site)

## Acknowledgements

We would like to thank Christina Myrdal, Bianca Marcellino (Dougan & Associates) and Genevieve Johnson for help collecting samples, and Carley Maitland for assisting with sample collection and processing. This study is funded by the government of Canada through Genome Canada and Ontario Genomics.

## Data Accessibility

Raw sequences will be available from NCBI SRA on acceptance. The bioinformatic pipeline SCVUC v4 is available from GitHub at https://github.com/Hajibabaei-Lab/SCVUC_COI_metabarcode_pipeline, the COI classifier v4 we used is available from https://github.com/terrimporter/CO1Classifier. The code used to generate figures, including infiles, are available from GitHub at https://github.com/terrimporter/RobinsonEtAl2020.

## Author Contributions and Conflicts of Interest

CVR, MTW, & MH designed the study, MTW conducted field work and laboratory analyses, TMP and CVR conducted statistical analyses and CVR wrote the manuscript with assistance from all of the authors.

## References

Baird, D. J., & Hajibabaei, M. (2012). Biomonitoring 2.0: a new paradigm in ecosystem assessment made possible by next-generation DNA sequencing. Molecular Ecology, 21(8), 2039–2044. doi: 10.1111/j.1365-294X.2012.05519.x

Barnes, M. A., & Turner, C. R. (2016). The ecology of environmental DNA and implications for conservation genetics. Conservation Genetics, 17(1), 1–17. doi: 10.1007/s10592-015-0775-4

Biggs, J., Ewald, N., Valentini, A., Gaboriaud, C., Dejean, T., Griffiths, R. A., … Dunn, F. (2015). Using eDNA to develop a national citizen science-based monitoring programme for the great crested newt (Triturus cristatus). Biological Conservation, 183, 19–28. doi:10.1016/j.biocon.2014.11.029

Bonada, N., Prat, N., Resh, V. H., & Statzner, B. (2006). DEVELOPMENTS IN AQUATIC INSECT BIOMONITORING: A Comparative Analysis of Recent Approaches. Annual Review of Entomology, 51(1), 495–523. doi:10.1146/annurev.ento.51.110104.151124

Brady, M. (2000). Alcohol Policy Issues for Indigenous People in the United States, Canada, Australia and New Zealand. Contemporary Drug Problems, 27(3), 435–509. doi: 10.1177/009145090002700304

Callahan, B. J., McMurdie, P. J., & Holmes, S. P. (2017). Exact sequence variants should replace operational taxonomic units in marker-gene data analysis. The ISME Journal, 11(12), 2639–2643. doi:10.1038/ismej.2017.119

Campbell, R. A. (2016). Making Sober Citizens: The Legacy of Indigenous Alcohol Regulation in Canada, 1777-1985. Journal of Canadian Studies. doi:10.3138/jcs.42.1.105

Demeke, T., & Jenkins, G. R. (2010). Influence of DNA extraction methods, PCR inhibitors and quantification methods on real-time PCR assay of biotechnology-derived traits. Analytical and Bioanalytical Chemistry, 396(6), 1977–1990. doi: 10.1007/s00216-009-3150-9

Dickie, I. A., Boyer, S., Buckley, H. L., Duncan, R. P., Gardner, P. P., Hogg, I. D., … Weaver, L. (2018). Towards robust and repeatable sampling methods in eDNA-based studies. Molecular Ecology Resources, 18(5), 940–952. doi: 10.1111/1755-0998.12907

Dixon, P. (2003). VEGAN, a package of R functions for community ecology. Journal of Vegetation Science, 14(6), 927–930. doi: 10.1111/j.1654-1103.2003.tb02228.x

Edgar, R. C. (2016). UNOISE2: improved error-correction for Illumina 16S and ITS amplicon sequencing. BioRxiv, 081257. doi:10.1101/081257

Emilson, C. E., Thompson, D. G., Venier, L. A., Porter, T. M., Swystun, T., Chartrand, D., … Hajibabaei, M. (2017). DNA metabarcoding and morphological macroinvertebrate metrics reveal the same changes in boreal watersheds across an environmental gradient. Scientific Reports, 7(1), 12777. doi: 10.1038/s41598-017-13157-x

Environment and Climate Change Canada. (2012). CABIN Wadeable Streams Manual. Retrieved from http://Publications.Gc.ca/Collections/Collection_2012/Ec/En84-87-2012-Eng.Pdf.

Geller, J., Meyer, C., Parker, M., & Hawk, H. (2013). Redesign of PCR primers for mitochondrial cytochrome c oxidase subunit I for marine invertebrates and application in all-taxa biotic surveys. Molecular Ecology Resources, 13(5), 851–861. doi: 10.1111/1755-0998.12138

Gibson, J., Shokralla, S., Porter, T. M., King, I., Konynenburg, S. van, Janzen, D. H., … Hajibabaei, M. (2014). Simultaneous assessment of the macrobiome and microbiome in a bulk sample of tropical arthropods through DNA metasystematics. Proceedings of the National Academy of Sciences, 111(22), 8007–8012. doi:10.1073/pnas.1406468111

Goldberg, C. S., Strickler, K. M., & Pilliod, D. S. (2015). Moving environmental DNA methods from concept to practice for monitoring aquatic macroorganisms. Biological Conservation, 183, 1–3. doi:10.1016/j.biocon.2014.11.040

Hajibabaei, M, Baird Donald J., Fahner Nicole A., Beiko Robert, & Golding G. Brian. (2016). A new way to contemplate Darwin’s tangled bank: how DNA barcodes are reconnecting biodiversity science and biomonitoring. Philosophical Transactions of the Royal Society B: Biological Sciences, 371(1702), 20150330. doi: 10.1098/rstb.2015.0330

Hajibabaei, Mehrdad, Porter, T. M., Robinson, C. V., Baird, D. J., Shokralla, S., & Wright, M. (2019). Watered-down biodiversity? A comparison of metabarcoding results from DNA extracted from matched water and bulk tissue biomonitoring samples. BioRxiv, 575928. doi:10.1101/575928

Hajibabaei, Mehrdad, Spall, J. L., Shokralla, S., & van Konynenburg, S. (2012). Assessing biodiversity of a freshwater benthic macroinvertebrate community through non-destructive environmental barcoding of DNA from preservative ethanol. BMC Ecology, 12(1), 28. doi: 10.1186/1472-6785-12-28

Hornung, B. V. H., Zwittink, R. D., & Kuijper, E. J. (2019). Issues and current standards of controls in microbiome research. FEMS Microbiology Ecology, 95(5). doi: 10.1093/femsec/fiz045

Kelling, S., Fink, D., La Sorte, F. A., Johnston, A., Bruns, N. E., & Hochachka, W. M. (2015). Taking a ‘Big Data’ approach to data quality in a citizen science project. Ambio, 44(4), 601–611. doi: 10.1007/s13280-015-0710-4

Kosmala, M., Wiggins, A., Swanson, A., & Simmons, B. (2016). Assessing data quality in citizen science. Frontiers in Ecology and the Environment, 14(10), 551–560. doi: 10.1002/fee.1436

Koziol, A., Stat, M., Simpson, T., Jarman, S., DiBattista, J. D., Harvey, E. S., … Bunce, M. (2019). Environmental DNA metabarcoding studies are critically affected by substrate selection. Molecular Ecology Resources, 19(2), 366–376. doi: 10.1111/1755-0998.12971

Ladell, B. A., Walleser, L. R., McCalla, S. G., Erickson, R. A., & Amberg, J. J. (2019). Ethanol and sodium acetate as a preservation method to delay degradation of environmental DNA. Retrieved from https://pubag.nal.usda.gov/catalog/6325757

Leray, M., Yang, J. Y., Meyer, C. P., Mills, S. C., Agudelo, N., Ranwez, V., … Machida, R. J. (2013). A new versatile primer set targeting a short fragment of the mitochondrial COI region for metabarcoding metazoan diversity: application for characterizing coral reef fish gut contents. Frontiers in Zoology, 10, 34. doi: 10.1186/1742-9994-10-34

Martin, M. (2011). Cutadapt removes adapter sequences from high-throughput sequencing reads. EMBnet.Journal, 17(1), 10–12. doi:10.14806/ej.17.1.200

McGee, K. M., Robinson, C. V., & Hajibabaei, M. (2019). Gaps in DNA-Based Biomonitoring Across the Globe. Frontiers in Ecology and Evolution, 7. doi: 10.3389/fevo.2019.00337

Minamoto, T., Naka, T., Moji, K., & Maruyama, A. (2016). Techniques for the practical collection of environmental DNA: filter selection, preservation, and extraction. Limnology, 17(1), 23–32. doi: 10.1007/s10201-015-0457-4

Muha, T. P., Rodríguez-Rey, M., Rolla, M., & Tricarico, E. (2017). Using Environmental DNA to Improve Species Distribution Models for Freshwater Invaders. Frontiers in Ecology and Evolution, 5. doi: 10.3389/fevo.2017.00158

NCBI Resource Coordinators. (2018). Database resources of the National Center for Biotechnology Information. Nucleic Acids Research, 46(D1), D8–D13. doi:10.1093/nar/gkx1095

Pawlowski, J., Kelly-Quinn, M., Altermatt, F., Apothéloz-Perret-Gentil, L., Beja, P., Boggero, A., … Kahlert, M. (2018). The future of biotic indices in the ecogenomic era: Integrating (e)DNA metabarcoding in biological assessment of aquatic ecosystems. Science of The Total Environment, 637-638, 1295–1310. doi: 10.1016/j.scitotenv.2018.05.002

Pedersen, Mikkel Winther, Overballe-Petersen, Søren, Ermini, Luca, Sarkissian, Clio Der, Haile, James, Hellstrom Micaela, … Willerslev Eske. (2015). Ancient and modern environmental DNA. Philosophical Transactions of the Royal Society B: Biological Sciences, 370(1660), 20130383. doi:10.1098/rstb.2013.0383

Pereira, H. M., & David Cooper, H. (2006). Towards the global monitoring of biodiversity change. Trends in Ecology & Evolution, 21(3), 123–129. doi:10.1016/j.tree.2005.10.015

Porter, T. M., & Hajibabaei, M. (2018). Automated high throughput animal CO1 metabarcode classification. Scientific Reports, 8(1), 4226. doi: 10.1038/s41598-018-22505-4

Rees, H. C., Maddison, B. C., Middleditch, D. J., Patmore, J. R. M., & Gough, K. C. (2014). REVIEW: The detection of aquatic animal species using environmental DNA - a review of eDNA as a survey tool in ecology. Journal of Applied Ecology, 51(5), 1450–1459. doi: 10.1111/1365-2664.12306

Rognes, T., Flouri, T., Nichols, B., Quince, C., & Mahé, F. (2016). VSEARCH: a versatile open source tool for metagenomics. PeerJ, 4, e2584. doi:10.7717/peerj.2584

RStudio Team. (2016). RStudio: Integrated Development Environment for R. Retrieved from http://Www.Rstudio.Com/.

Sales, N. G., Wangensteen, O. S., Carvalho, D. C., & Mariani, S. (n.d.). Influence of preservation methods, sample medium and sampling time on eDNA recovery in a neotropical river. Environmental DNA, 0(0). doi:10.1002/edn3.14

Schrader, C., Schielke, A., Ellerbroek, L., & Johne, R. (2012). PCR inhibitors - occurrence, properties and removal. Journal of Applied Microbiology, 113(5), 1014–1026. doi:10.1111/j.1365-2672.2012.05384.x

Shapiro, S. S., & Wilk, M. B. (1965). An Analysis of Variance Test for Normality (Complete Samples). Biometrika, 52(3/4), 591–611. doi:10.2307/2333709

St. John, J. (2016). SeqPrep. Retrieved from https://Github.Com/Jstjohn/SeqPrep/Releases.

Steininger, S., Storer, C., Hulcr, J., & Lucky, A. (2015). Alternative preservatives of insect DNA for citizen science and other low-cost applications. Invertebrate Systematics, 29(5), 468–472. doi: 10.1071/IS15003

Strickler, K. M., Fremier, A. K., & Goldberg, C. S. (2015). Quantifying effects of UV-B, temperature, and pH on eDNA degradation in aquatic microcosms. Biological Conservation, 183, 85–92. doi:10.1016/j.biocon.2014.11.038

Taberlet, P., Bonin, A., Zinger, L., & Coissac, E. (2018). Environmental DNA: For Biodiversity Research and Monitoring. Oxford University Press.

Taberlet, P., Coissac, E., Hajibabaei, M., & Rieseberg, L. H. (2012). Environmental DNA. Molecular Ecology, 21(8), 1789–1793. doi:10.1111/j.1365-294X.2012.05542.x

Thomsen, P. F., & Willerslev, E. (2015). Environmental DNA - An emerging tool in conservation for monitoring past and present biodiversity. Biological Conservation, 183, 4–18. doi:10.1016/j.biocon.2014.11.019

Turner, C. R., Uy, K. L., & Everhart, R. C. (2015). Fish environmental DNA is more concentrated in aquatic sediments than surface water. Biological Conservation, 183, 93–102. doi:10.1016/j.biocon.2014.11.017

Vasselon, V., Domaizon, I., Rimet, F., Kahlert, M., & Bouchez, A. (2017). Application of high-throughput sequencing (HTS) metabarcoding to diatom biomonitoring: Do DNA extraction methods matter? Freshwater Science, 36(1), 162–177. doi: 10.1086/690649

Wang, Q., Garrity, G. M., Tiedje, J. M., & Cole, J. R. (2007). Naïve Bayesian Classifier for Rapid Assignment of rRNA Sequences into the New Bacterial Taxonomy. Applied and Environmental Microbiology, 73(16), 5261–5267. doi:10.1128/AEM.00062-07

Williams, K. E., Huyvaert, K. P., & Piaggio, A. J. (2016). No filters, no fridges: a method for preservation of water samples for eDNA analysis. BMC Research Notes, 9(1), 298. doi: 10.1186/s13104-016-2104-5

Williams, S. T. (2007). Safe and legal shipment of tissue samples: does it affect DNA quality? Journal of Molluscan Studies, 73(4), 416–418. doi: 10.1093/mollus/eym039

WWF-Canada. (2017). A national assessment of Canada’s freshwater: Watershed Reports, 3–23.

Yamanaka, H., Minamoto, T., Matsuura, J., Sakurai, S., Tsuji, S., Motozawa, H., … Kondo, A. (2017). A simple method for preserving environmental DNA in water samples at ambient temperature by addition of cationic surfactant. Limnology, 18(2), 233–241. doi: 10.1007/s10201-016-0508-5

Yang, J., Jeppe, K., Pettigrove, V., & Zhang, X. (2018). Environmental DNA Metabarcoding Supporting Community Assessment of Environmental Stressors in a Field-Based Sediment Microcosm Study. Environmental Science & Technology, 52(24), 14469–14479. doi:10.1021/acs.est.8b04903

Zinger, L., Bonin, A., Alsos, I. G., Bálint, M., Bik, H., Boyer, F., … Taberlet, P. (2019). DNA metabarcoding–Need for robust experimental designs to draw sound ecological conclusions. Molecular Ecology, 28(8), 1857–1862. doi:10.1111/mec.15060

