## Supplementary Material for "Propylene glycol-based antifreeze as an effective preservative for DNA metabarcoding of benthic arthropods"

**Table S1.** Information on sites kick net sampled for benthos, including site number, site name, latitude, longitude and number of samples collected.

| Site number | Site name | Latitude | Longitude | Number of samples collected |
| --- | --- | --- | --- | --- |
| 1 | Laurel 10-3 | 43.4841207 | -80.566787 | 2 |
| 2 | Laurel 7-2 | 43.4707269 | -80.556274 | 2 |
| 3 | Laurel 4-3 | 43.4676369 | -80.5332779 | 2 |
| 4 | Beaver 19 | 43.4911767 | -80.624394 | 1 |
| 5 | Beaver 18 | 43.4920012 | -80.609783 | 1 |
| 6 | Clair 12 | 43.46540880 | -80.571321 | 1 |

**Table S2. Summary of read processing**

| Step | BR5 | F230R | MiCOI | Total |
| --- | --- | --- | --- | --- |
| Raw | - | - | - | 11,185,532 x 2 |
| Paired | - | - | - | 10,767,127 |
| Primer trimmed | 2,837,670 | 1,980,177 | 5,090,979 | 9,908,826 |

**Table S3. Summary of ESV processing**

| Step | BR5 | F230R | MiCOI | Total |
| --- | --- | --- | --- | --- |
| Denoised* ESVs | 10,868 | 4,719 | 9,397 | 24,984 |
| Denoised ESVs, putative pseudogenes removed | 7,883 | 4,122 | 7,293 | 19,298 |
| Arthropoda ESVs | 1,711 | 1,320 | 2,167 | 5,198 |
| Reads in Arthropoda ESVs | 683,908 | 666,732 | 1,067,812 | 2,418,452 |

*Denoising refers to removal of ESVs with putative sequence errors, PhiX contamination, rare sequences, and putative chimeric sequences

**Table S4. COI order data for the negative controls amplified in this study.** The COI data set was filtered for bacteria, fungi, plants, algae, mammals, fish, and protists.

| Order | Antifreeze | Ethanol | Paired Extraction Kit | Paired PCR | Single Extraction Kit | Single PCR |
| --- | --- | --- | --- | --- | --- | --- |
| Araneae | 0 | 0 | 0 | 3 | 0 | 0 |
| Coleoptera | 0 | 0 | 3 | 0 | 0 | 0 |
| Diptera | 3 | 0 | 21 | 21 | 16 | 0 |
| Entomobryomorpha | 0 | 0 | 0 | 0 | 11 | 0 |
| Ephemeroptera | 15 | 0 | 3 | 9 | 49 | 0 |
| Euryalida | 4 | 12 | 7 | 6 | 9 | 0 |
| Haplotaxida | 0 | 80 | 7 | 9 | 9 | 0 |
| Hemiptera | 0 | 0 | 4 | 4 | 3 | 0 |
| Hirudinida | 0 | 0 | 0 | 0 | 4 | 0 |
| Isopoda | 3 | 0 | 0 | 0 | 0 | 0 |
| Lepidoptera | 0 | 9 | 0 | 0 | 0 | 0 |
| Trichoptera | 6 | 0 | 11 | 5 | 6 | 68 |
| Trombidiformes | 0 | 0 | 0 | 8 | 0 | 0 |

**Figure S1.** Map of sampling locations for single ‘mock’ samples (Beaver 19, Beaver 18 and Clair 12) and paired samples (Laurel 10-3, Laurel 7-2 and Laurel 4-3), located within tributaries of the Grand River (Waterloo, ON).

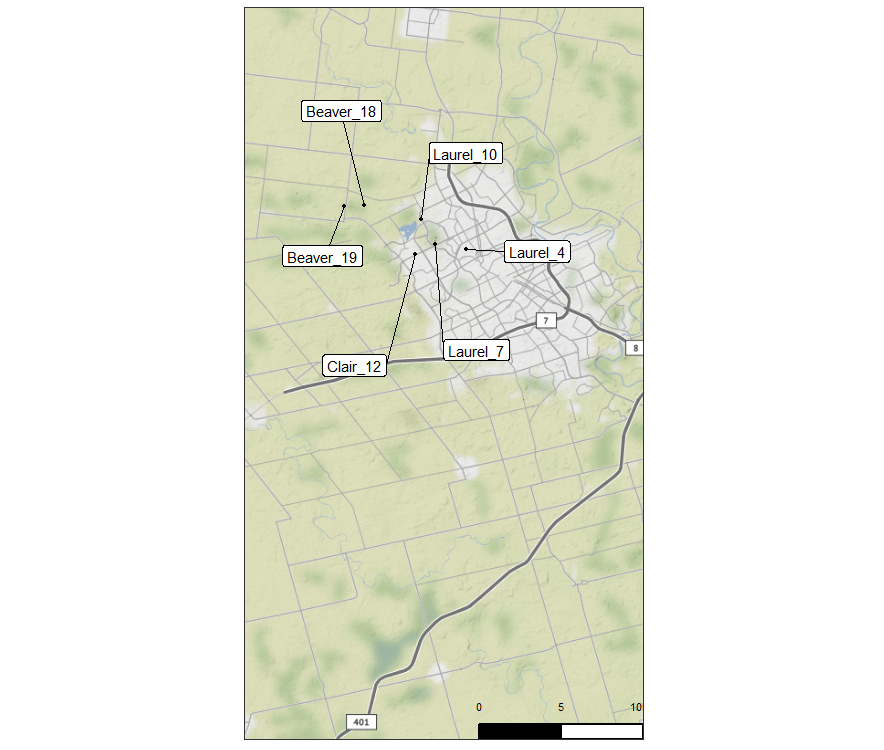

**Figure S2. Antifreeze samples produced similar quality bands to ethanol samples across the three primer sets. Gel shows bands for paired samples only.** For each site (1-3): A# = ethanol samples processed using method 2; E# = ethanol samples processed using method 2; a# = antifreeze samples processed using method 1; e# = ethanol samples processed using method 1. Negative controls: a- = antifreeze blank; et- = ethanol blank; ex- = extraction kit blank; p1 = PCR negative control 1 (carried over from first PCR); p2 = PCR negative control 2 (for second PCR only).

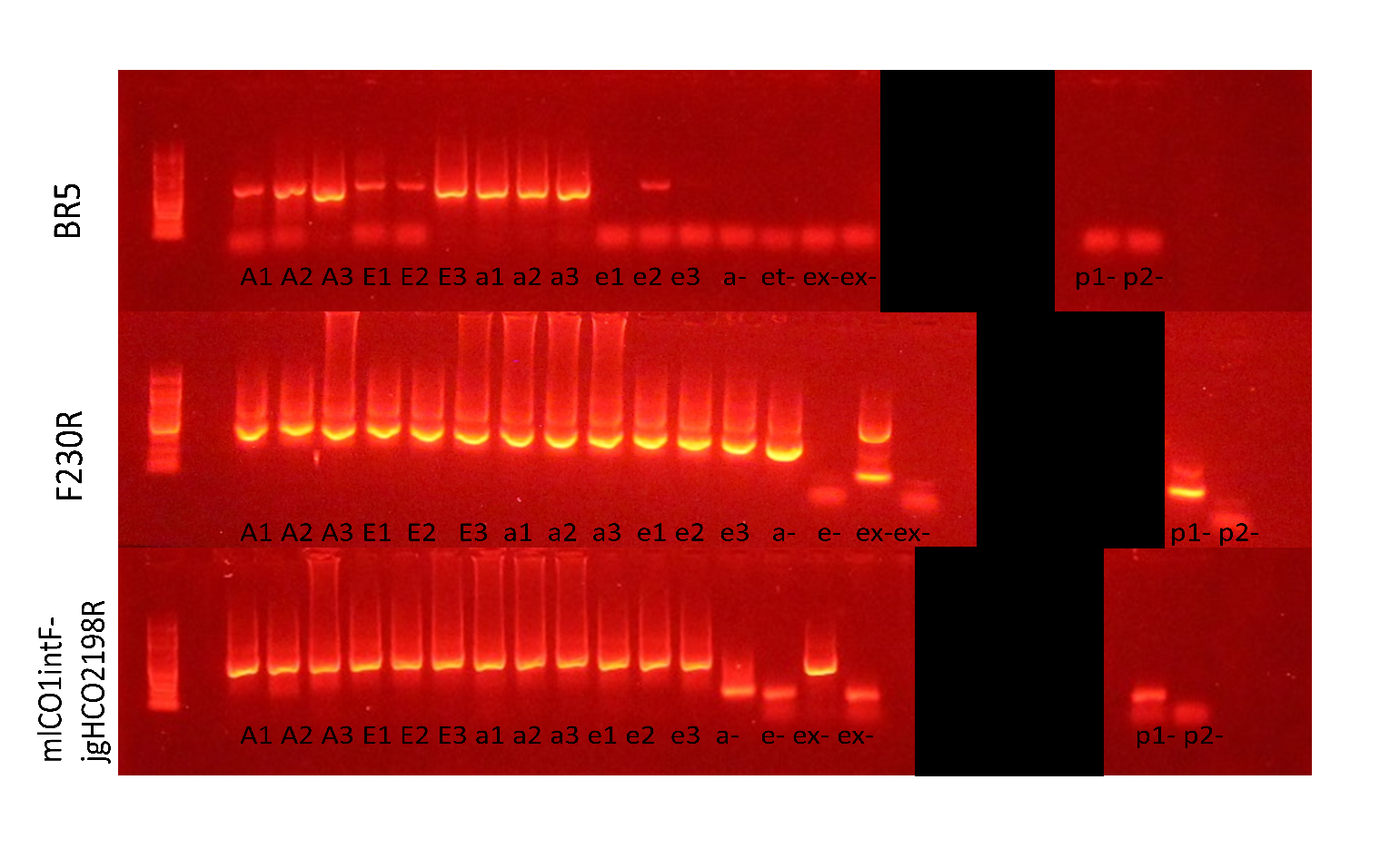

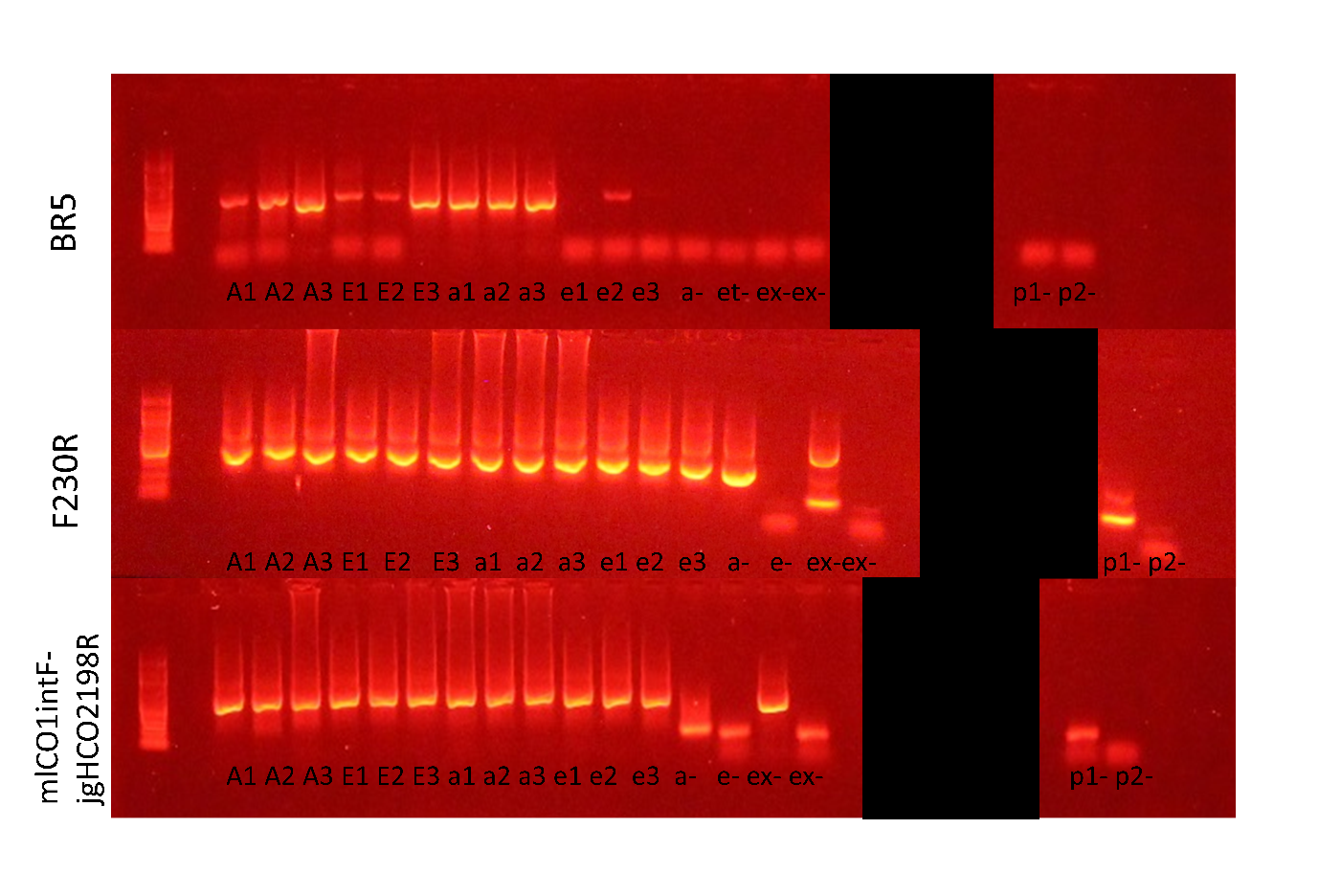

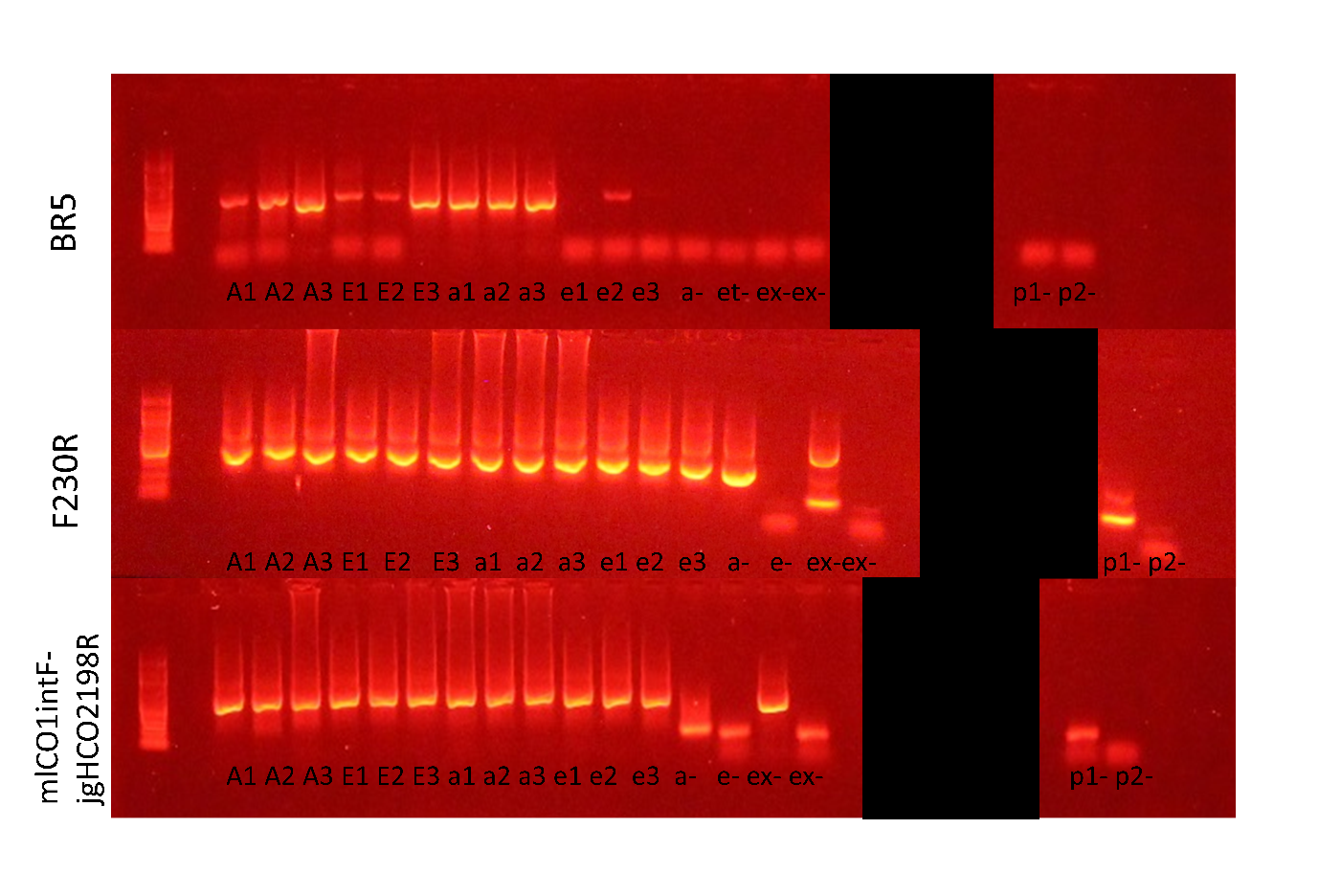

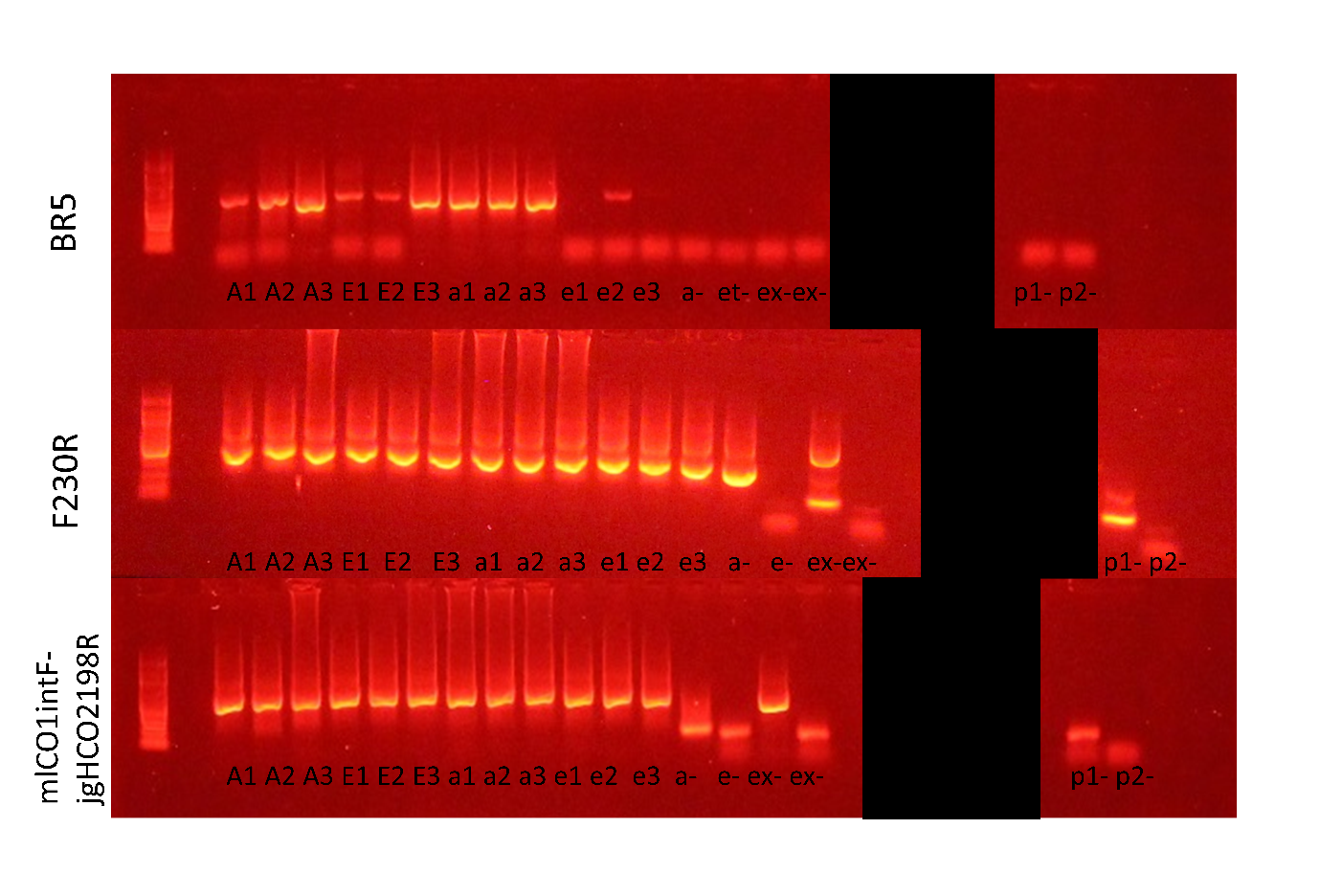

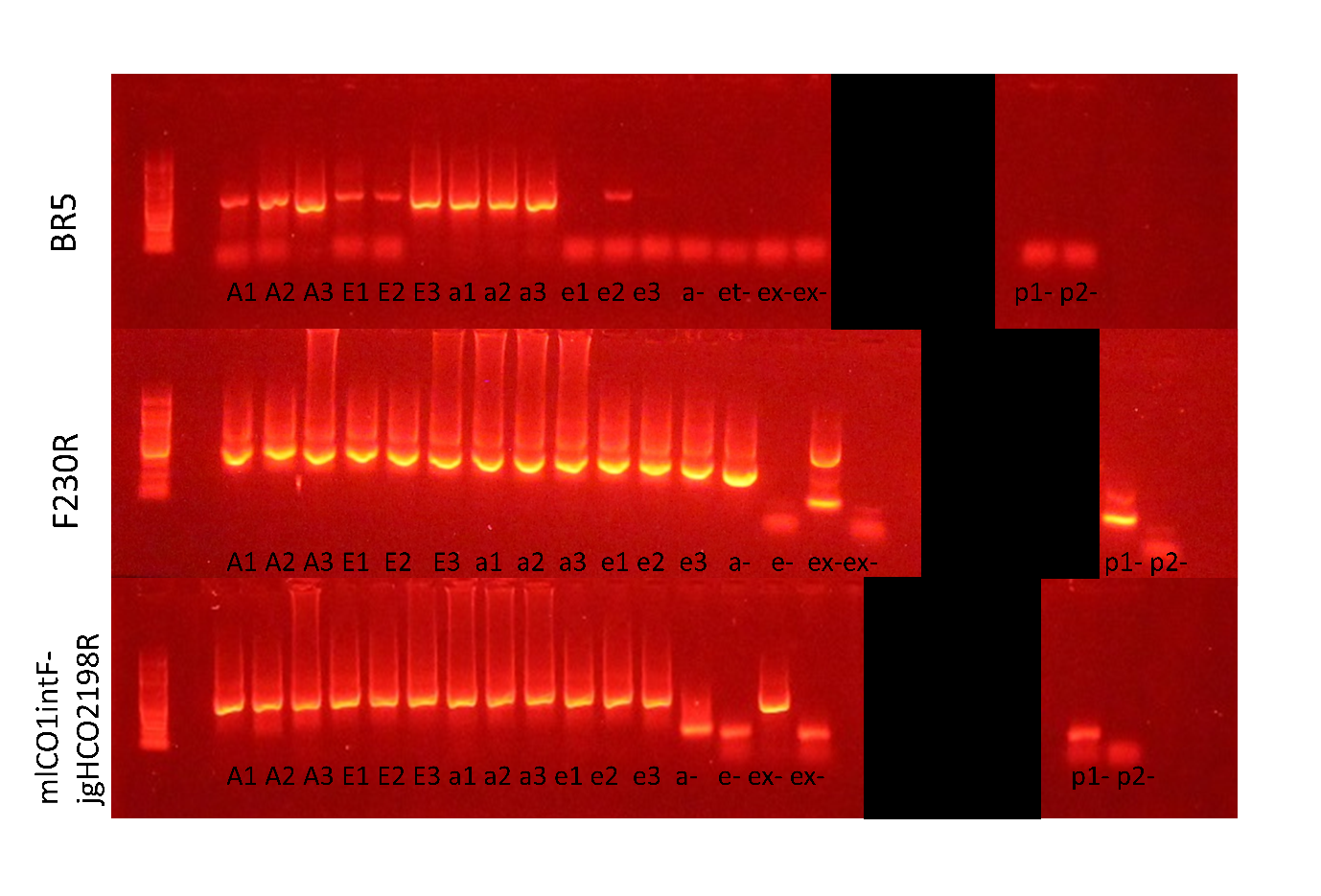

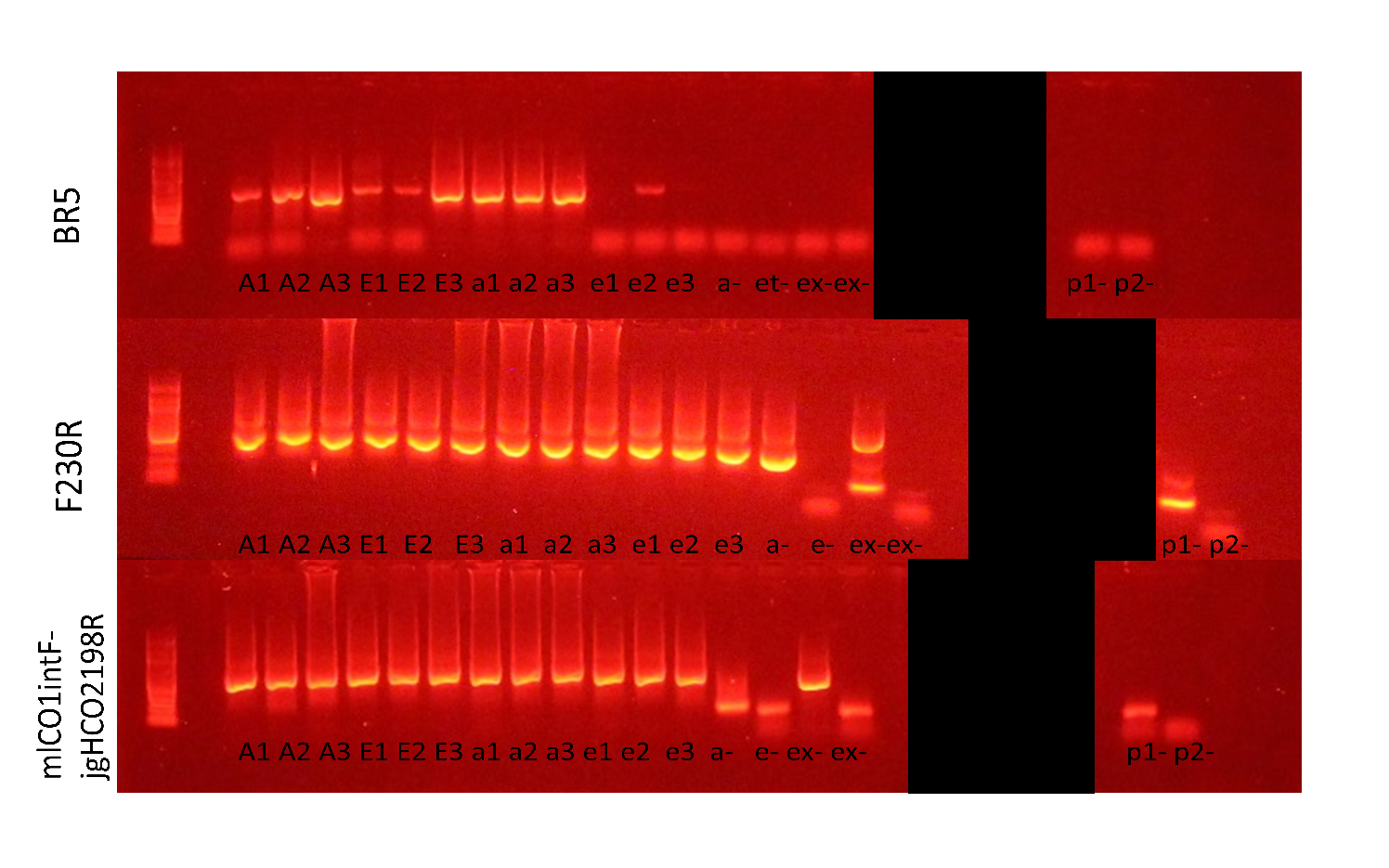

**Figure S3. Arthropoda is the most abundant phylum recovered.** Results for the top 10 phyla are shown with respect to proportion of ESVs and reads recovered. Based on raw unnormalized data.

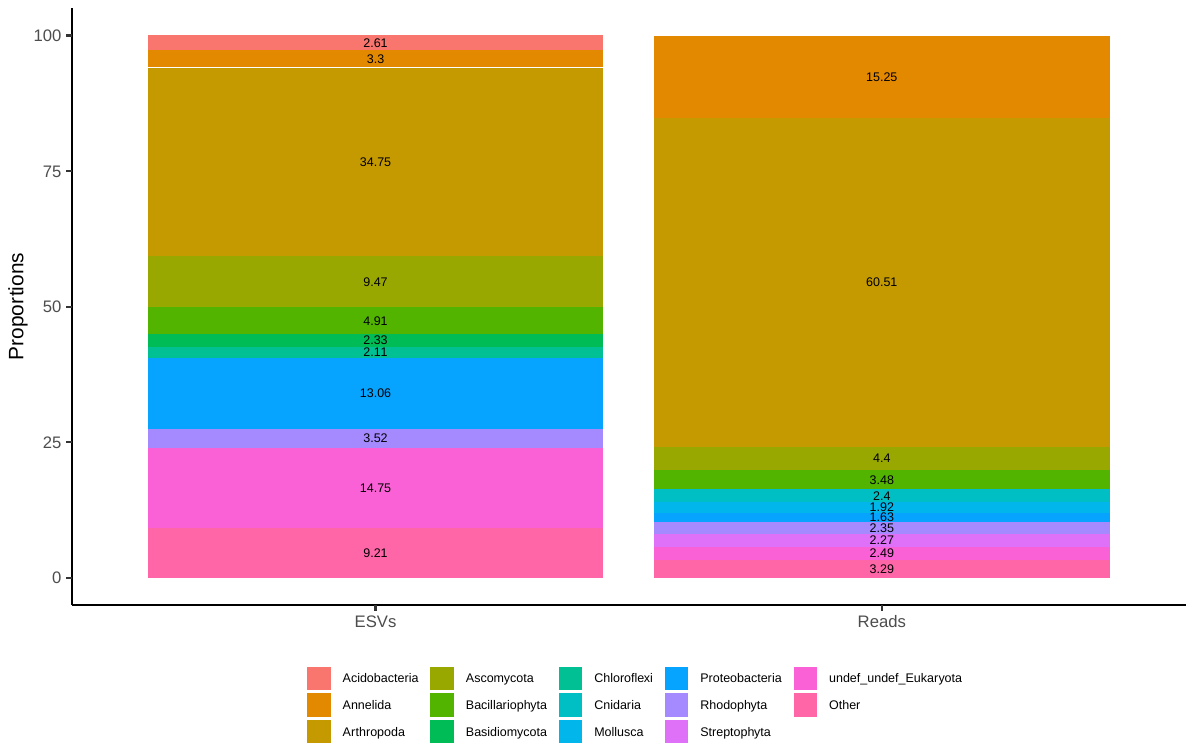

**Figure S4. All samples show that ESV sampling reached saturation.** The top row shows curves based on all ESVs; the bottom row shows curves based on Arthropoda ESVs only. Samples were color-coded by site or treatment as shown in the legend. The vertical dashed line indicates the 15^th^ percentile of sampling read depth (excluding controls), which is the number of reads that would be used in any future analysis based on normalized data.

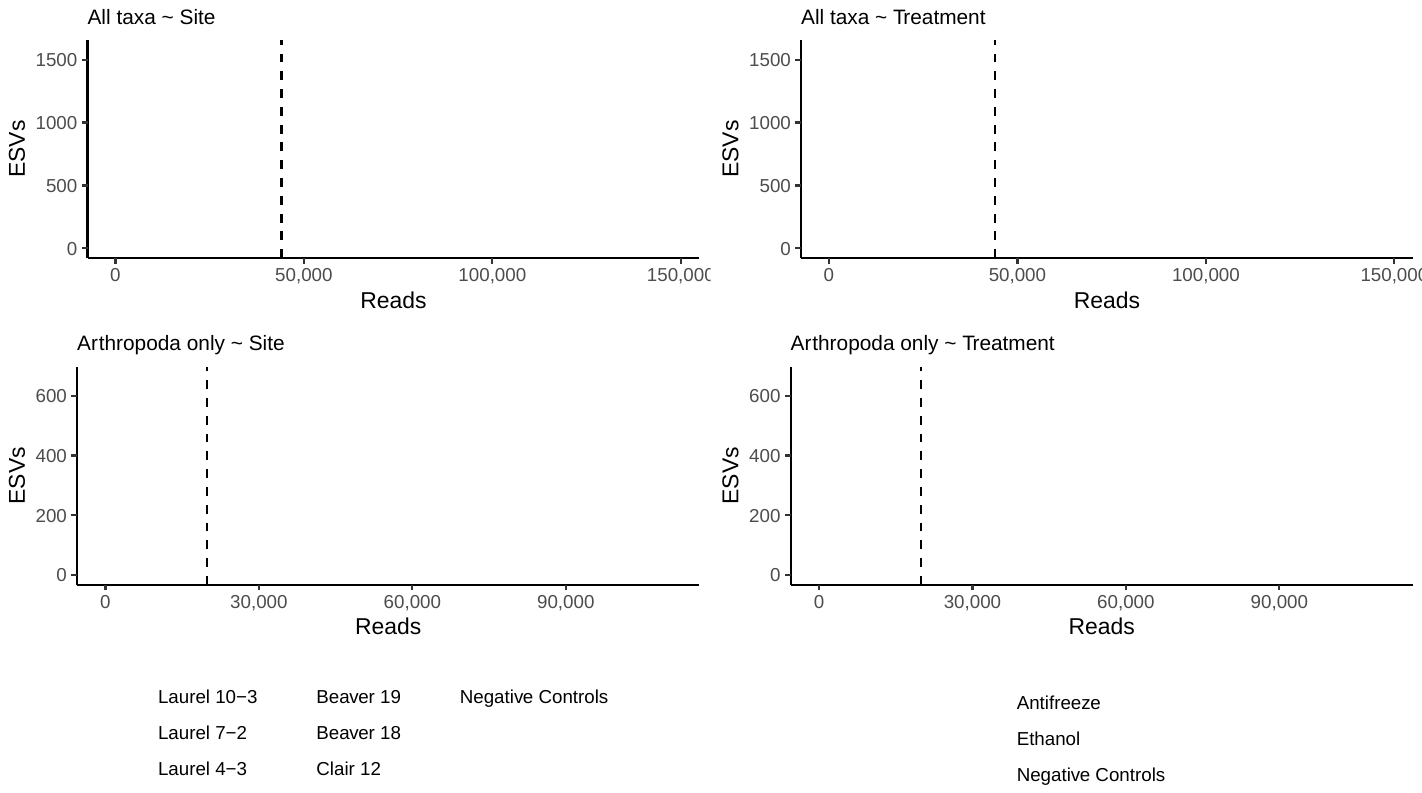

**Figure S5. Many arthropod species and most arthropod genera and families were confidently identified using three different COI amplicons.** Results based on raw unnormalized data.

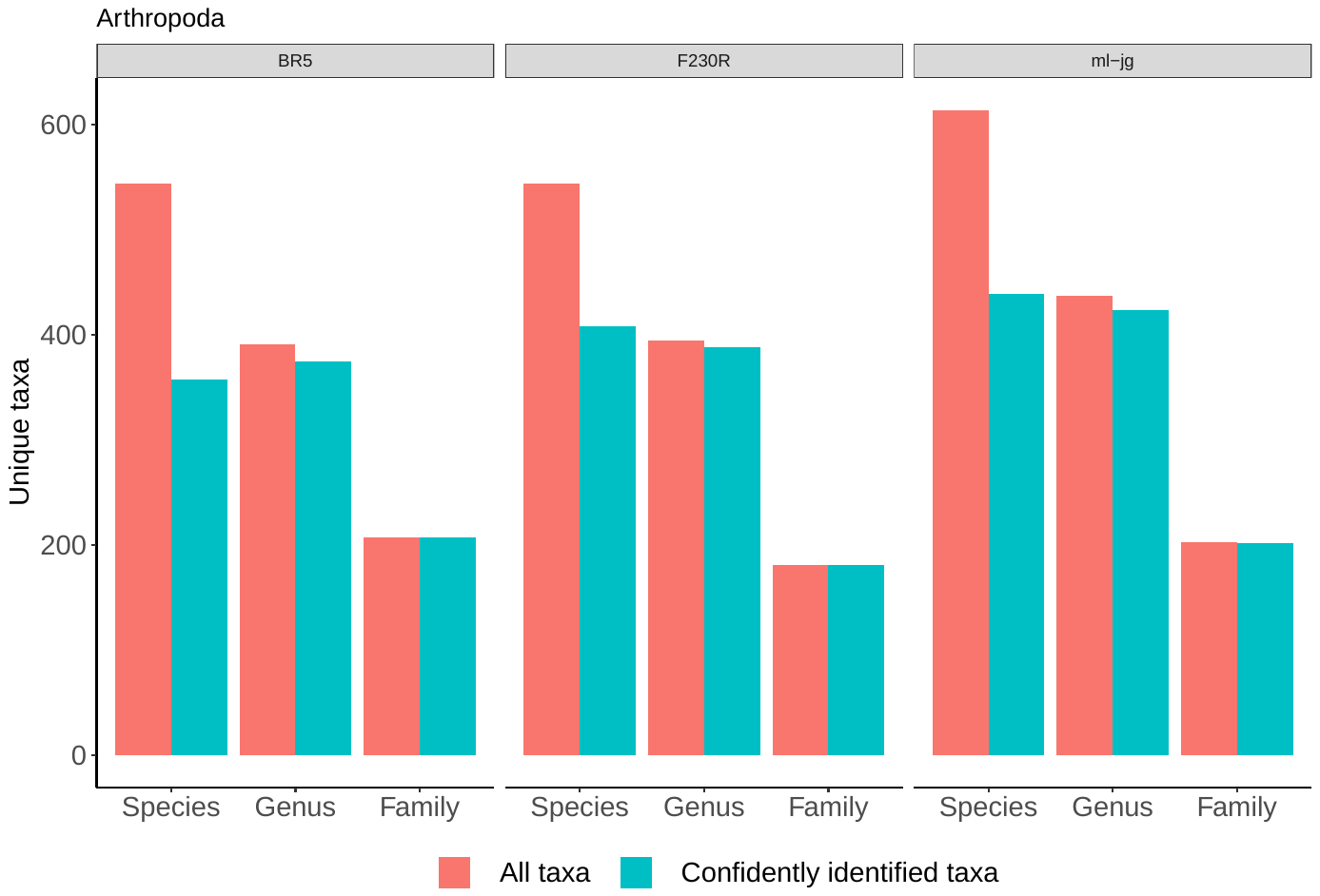
